# Structural characterization of tau in fuzzy tau:tubulin complexes

**DOI:** 10.1101/647016

**Authors:** Ho Yee Joyce Fung, Kristen McKibben, Jennifer Ramirez, Kushol Gupta, Elizabeth Rhoades

**Affiliations:** Department of Chemistry, University of Pennsylvania, Philadelphia, PA 19104, USA; Biochemistry and Molecular Biophysics Graduate Group, Perelman School of Medicine at University of Pennsylvania, Philadelphia, PA 19104, USA; Department of Biochemistry and Biophysics, Perelman School of Medicine at University of Pennsylvania, Philadelphia, PA 19104, USA

**Keywords:** tau, tubulin, microtubule, Intrinsically disordered protein, acrylodan, fluorescence correlation spectroscopy

## Abstract

Tau is a neuronal microtubule (MT) associated protein of significant interest due to its association with several neurodegenerative disorders. Tau’s intrinsic disorder and the dynamic nature of its interactions with tubulin and MTs make its structural characterization challenging. Here we use an environmentally sensitive fluorophore as a site-specific probe of tau bound to soluble tubulin. Comparison of our results with recently published tau:MT cryo-EM model reveals structural similarities between tubulin- and MT-bound tau. Analysis of residues across the repeat regions reveal a hierarchy in tubulin occupancy, which may be relevant to tau’s ability to differentiate between tubulin and MTs. As binding to soluble tubulin is a critical first step in MT polymerization, our characterization of the structural features of tau in dynamic, fuzzy tau:tubulin assemblies advances our understanding of how tau functions in the cell and how function may be disrupted in disease.

## Introduction

Tau is a predominantly neuronal protein that is the focus of significant attention due to a proposed causal role in a variety of neurodegenerative disorders, including Alzheimer’s disease (Gao et al., 2018). Natively tau is a microtubule (MT) associated protein with a putative role in regulating the dynamics of axonal MTs (Drubin and Kirschner, 1986; Weingarten et al., 1975). Tau is both developmentally regulated by alternative splicing and post-translationally regulated to modulate its interaction with MTs, and dysregulation of these processes can lead to pathological aggregation of tau (Kosik et al., 1989; Martin et al., 2011; Park et al., 2016). Despite extensive research on the physiological and pathological roles of tau, details of the mechanisms of normal tau function in MT polymerization remains unclear.

Tau is an intrinsically disordered protein that lacks stable secondary and tertiary structure (Cleveland et al., 1977; Woody et al., 1983). It has multiple binding sites for tubulin distributed throughout its proline-rich region (PRR), four MT binding repeats (MTBR: R1 – R4) and pseudo-repeat (R’) (Figure 1A) (Butner and Kirschner, 1991; Fauquant et al., 2011; Gustke et al., 1994; McKibben and Rhoades, 2019; Mukrasch et al., 2007). When tubulin is present in high excess of tau, heterogenous tau:tubulin complexes are formed, and the ability to bind multiple tubulin dimers is key to tau’s role in polymerization of MTs (Li and Rhoades, 2017). The dynamic nature of tau and tau:tubulin interactions have impeded structure-function studies and despite intensive study, much is still unclear about how tau binds MTs and soluble tubulin.

**Figure 1.**
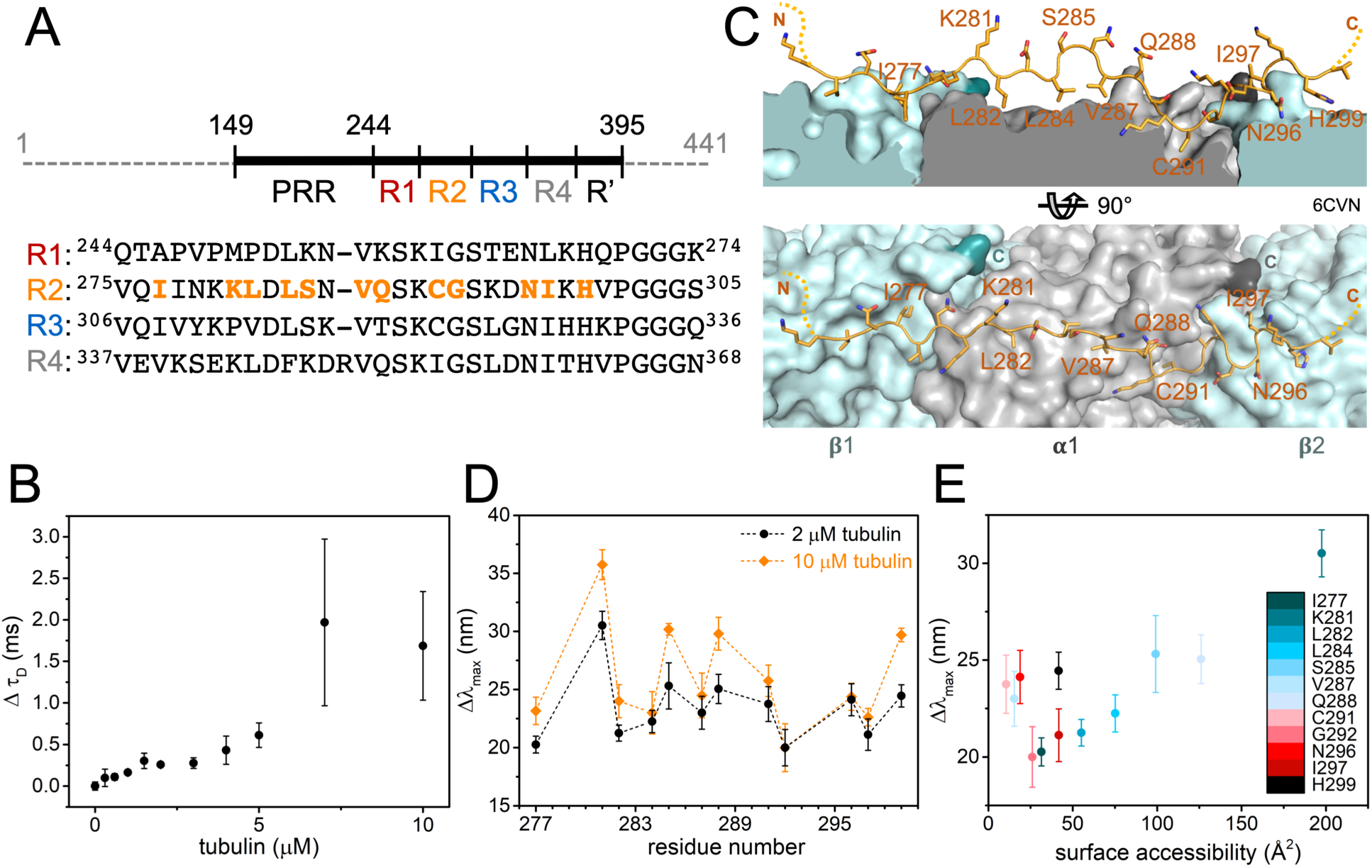
Structure of tubulin- and MT-bound tau are similar. (A) Schematic of tau construct (residues 149 – 395) with domains labeled: proline-rich region (PRR), MT-binding repeats (R1 – R4) and pseudo repeat (R’). Sequence alignment of R1 – R4 is shown below with positions screened in this study highlighted in orange. (B) Change in diffusion time (Δτ_D_) of 300 nM tau_A488_ upon tubulin binding. Data points are mean ± SD, n=3. See also Figure S1. (C) Structural model of tau:MT (PDB ID:6CVN) where tau is shown as orange cartoon and α/β-tubulin as grey/light blue surfaces with their C-terminal residues highlighted in dark grey and turquoise respectively. Disordered C-terminal tubulin tails are not resolved in the model. (D) Magnitude of emission blue-shift (Δλ_max_) of tau_acrylodan_ at different positions in R2 in the presence of 2 μM or 10 μM tubulin. Data points are mean ± SD, n≥3. See also Figure S2. (E) Δλ_max_ in the presence of 2 μM tubulin plotted against surface accessibility of respective residues in tau:MT structure calculated using GetArea (Fraczkiewicz and Braun, 1998). Data points are color-coded by residue position.

Numerous studies have identified putative binding sites of tau on soluble tubulin and the MT lattice, as well as characterized structural features of bound tau (Al-Bassam et al., 2002; Chau et al., 1998; Kadavath et al., 2018; Kadavath et al., 2015; Kar et al., 2003; Martinho et al., 2018; Santarella et al., 2004; Serrano et al., 1985). Most recently, cryo-EM and computational modeling revealed the first near-atomic level model of MT-bound tau (Kellogg et al., 2018). Tandem repeats of R1 and R2 were used to improve resolution of the cryo-EM map, which showed segments of the repeats decorating the outer surface of each MT protofilament. Each tau repeat is shown to adopt an overall extended conformation, spanning across two tubulin heterodimers with a high affinity binding site at the interdimer interface and weak interactions over the intradimer surface.

Prior work from our lab using acrylodan, an environmentally sensitive fluorophore, to map the structural features of R3 in tubulin-bound tau observed a periodic pattern that we proposed was consistent with helical structure (Li et al., 2015). The extended conformation of tau in the cryo-EM model motivated us to re-examine our interpretation of this data and to investigate possible structural differences between tubulin- and MT-bound tau. Here, in order to make a more direct comparison with the EM work, we use acrylodan fluorescence to profile the structural features of R2 in tubulin-bound tau and to test for hierarchical binding sites in the other repeats. Overall, our work provides details of the structural features of tau in heterogenous ‘fuzzy’ tau-tubulin complexes, which provide insight for possible mechanisms in tau-mediated MT polymerization.

## Results

### Structural features of tubulin- and MT-bound tau are similar

For this work, we used fragment of tau (residues 149-395) consisting of all of the regions implicated in direct interactions with soluble tubulin or MTs: PRR, MTBR and R’ (Figure 1A). Fluorescence correlation spectroscopy (FCS) was first used to quantify tau binding to tubulin. Titration of tubulin into 300 nM tau labeled with Alexa Fluor 488 (tau_A488_) revealed a biphasic binding curve where the first saturation was achieved at around 2 μM tubulin, corresponding to an increase in diffusion time (τ_D_) of ∼0.25 ms relative to tau alone (Figure 1B). Analysis of individual autocorrelation curves and counts per molecule (CPM) showed that this first saturation consists of one tau molecule (Figure S1A and 1B). The magnitude of the increase in τ_D_, and supporting measurements by analytical ultracentrifugation (Figure S1C – H), is consistent with a single tau bound to two tubulin dimers (Li et al., 2015). A 1:2 tau:tubulin dimer stoichiometry has been reported for MTs by several different groups (Fauquant et al., 2011; Gustke et al., 1994). Further increasing the tubulin concentration caused large increases in both τ_D_ and CPM, indicative of heterogeneous, non-covalently crosslinked tau:tubulin complexes (Figure S1A and S1B), described previously (Li and Rhoades, 2017). Based on these results, we decided to test the response of acrylodan-labeled tau at the two saturating concentrations, 2 μM and 10 μM tubulin, to probe the structural features of R2 in both small and large tau:tubulin complexes.

Cysteine mutations were introduced individually throughout R2 to allow for site-specific labeling with acrylodan (Figure 1A), in order to compare with our previous work with R3 (Li et al., 2015), as well as the published tau:MT structure (Figure 1C) (Kellogg et al., 2018). Anisotropy measurements comparing tau labeled with acrylodan (tau_acrylodan_) at two different sites within R2 with tau_A488_ labeled at the N-terminus show that labeling with acrylodan in the MTBR does not significantly impair tubulin binding (Figure S2A). The fluorescence emission spectra of tau_acrylodan_ were measured in the absence and presence of 2 μM or 10 μM tubulin and the emission maximum (λ_max_) was quantified. In the absence of tubulin, λ_max_ was independent of labeling position (Figure S2B). Binding of tau to tubulin resulted in a shift of the emission spectrum to shorter wavelengths, i.e. a blue-shift, as expected (Hibbs et al., 2004; Stuart et al., 2011; Sun et al., 2007). This is due to a reduction in polarity of acrylodan’s surrounding environment upon binding (Figure S2C) (Prendergast et al., 1983). All labeling positions tested exhibited emission blue-shifting upon binding tubulin, although there was significant site-dependent and tubulin concentration-dependent variability in the magnitudes of the shifts (Figure 1D). As tau:tubulin interactions are highly dynamic, both the local environment of the acrylodan when bound as well as the average occupancy of any binding site will impact the magnitude of the blue-shift.

For both concentrations of tubulin, plots of the peak shift magnitudes (Δλ_max_) as a function of R2 sequence showed a periodic pattern (Figure 1D). The overall shape and periodicity of the Δλ_max_ profiles are the same at both tubulin concentrations, suggesting that instead of a drastic reorganization of the R2 structure, further acrylodan peak shifts at 10 μM likely reflect increased occupancy of tubulin binding sites in R2. The general pattern of the Δλ_max_ magnitudes observed for R2 residues 281 – 291 is in good agreement with that previously reported for the equivalent residues in R3 (312 – 322) (Figure S2D) (Li et al., 2015); specifically, both R2 and R3 show peaks and troughs in Δλ_max_ at equivalent residues. This data supports similar tubulin-binding modes between the different repeats for these residues. However, the pattern diverges for residues 292 – 297 in R2 from that observed for residues 323 – 328 in R3 (Figure S2D). These residues are found at the interdimer interface in the tau:MT EM structure (Figure 1C), an interface that is not stably present for soluble tubulin. Differences in Δλ_max_ in this region between the R2 and R3 may reflect the extent to which binding sites in their neighboring repeats are populated, i.e. whether R3 is populated for R2 and whether R4 is populated for R3. Moreover, binding to two tubulin dimers to adjacent repeat regions may effectively impose formation of an interdimer interface. Interestingly, the Δλ_max_ of residues in R2 closest to the interdimer interface or “anchor point” on MTs (residues G292 and N296) saturated at 2 μM tubulin, while the remainder of the positions tested showed varying additional increases at 10 μM tubulin (Figure 1C and 1D). This is consistent with the observation based on the cryo-EM structure that tau has a higher occupancy at the interdimer interface relative to the intradimer surface (Kellogg et al., 2018).

A more detailed comparison between our measurements and the EM structure was carried out to provide context for our results. We find an overall positive correlation between Δλ_max_ and the surface accessibility of the residues determined from the tau:MT structural model (Figure 1E). This observation is initially non-intuitive, as residues that are more shielded from the surrounding solvent upon binding to tubulin are expected to experience a less-polar environment and thus larger blue-shifts in the acrylodan emission spectra at these residues. However, the excited state of the conjugated acrylodan has a positive dipole oriented away from the cysteine sidechain which can interact with the charges in its surrounding, causing a reduction in blue-shift of its emission (Cerezo et al., 2001). There are an abundance of negative charges on the surface of tubulin, making it plausible that acrylodan conjugated at residues that are oriented towards the tubulin surface interact with surface residues in a manner that results in a reduction in Δλ_max_.

There are several notable deviations from the positive correlation, i.e. residues with relatively low surface accessibility but large blue-shifts in their emission spectra. Residues V287 and C291 have buried binding sites on α-tubulin, with the least accessible surface area (<25 Å^2^) of the residues tested (Figure 1E). One possibility is that acrylodan is well-buried in the binding pockets and exhibits the expected large blue-shift upon shielding from solvent. Alternatively, the acrylodan label may not fit well into the binding pockets, causing the fluorophore to be partially excluded from the tubulin surface and unable to form blue-shift reducing interactions as described above (Figure 1E). Residues N296 and H299, on the other hand, bind to the adjacent tubulin dimer in MTs (Figure 1C). The lack of a stable interdimer interface in soluble tubulin may mean that these residues interact with tubulin surfaces that are buried and therefore inaccessible in MTs. However, if tau is able to impose the interdimer interface, as we suggest above, the relatively large shifts in these residues may reflect the increased occupancy at this interface. It is also possible that acrylodan is not equally well tolerated at all positions because it disrupts interactions with tubulin specific to the amino acid it replaces.

### Disordered tubulin tails enhance tau binding but minimally impact R2 structure

The cryo-EM structure suggests that tau’s polar residues point away from the tubulin surface where they may be poised to interact with the acidic tubulin tails, although these are not resolved in the structure (Kellogg et al., 2018). In order to test the possible interaction between tubulin tails and tau, we generated tailless tubulin by subtilisin cleavage (Figure S3A) and tested the effect on tau binding and acrylodan peak shifts. FCS was used to compare binding of tau to intact and tailless tubulin. Titration of tubulin into 20 nM tau_A488_ revealed that tau binds tailless tubulin with approximately five-fold weaker affinity that that of intact tubulin (Figure 2A). Treatment control of tubulin without the addition of protease showed similar binding to tau when compared to untreated tubulin, indicating that the treatment itself does not cause a reduction in tau binding (Figure 2A). The maximum increase in τ_D_ of tau upon binding tailless tubulin is ∼0.15 ms, smaller than the increase seen for intact tubulin at comparable concentrations (Figure 1B, 2A and S3B), consistent both with the weaker affinity and a reduced tau:tubulin stoichiometry. Moreover, binding of 300 nM tau_A488_ to tailless tubulin did not show any evidence of the biphasic binding (Figure S3B) nor formation of large tau-tubulin assemblies (Figure S3C and S3D), in striking contrast to our observations with intact tubulin (Figures 1B, S1A and S1B). Based on the FCS analysis, 4 μM tailless tubulin, corresponding to the plateau in the increase in τ_D_ (Figure S3B), was selected for acrylodan measurements for comparison with the 2 μM intact tubulin measurements.

**Figure 2.**
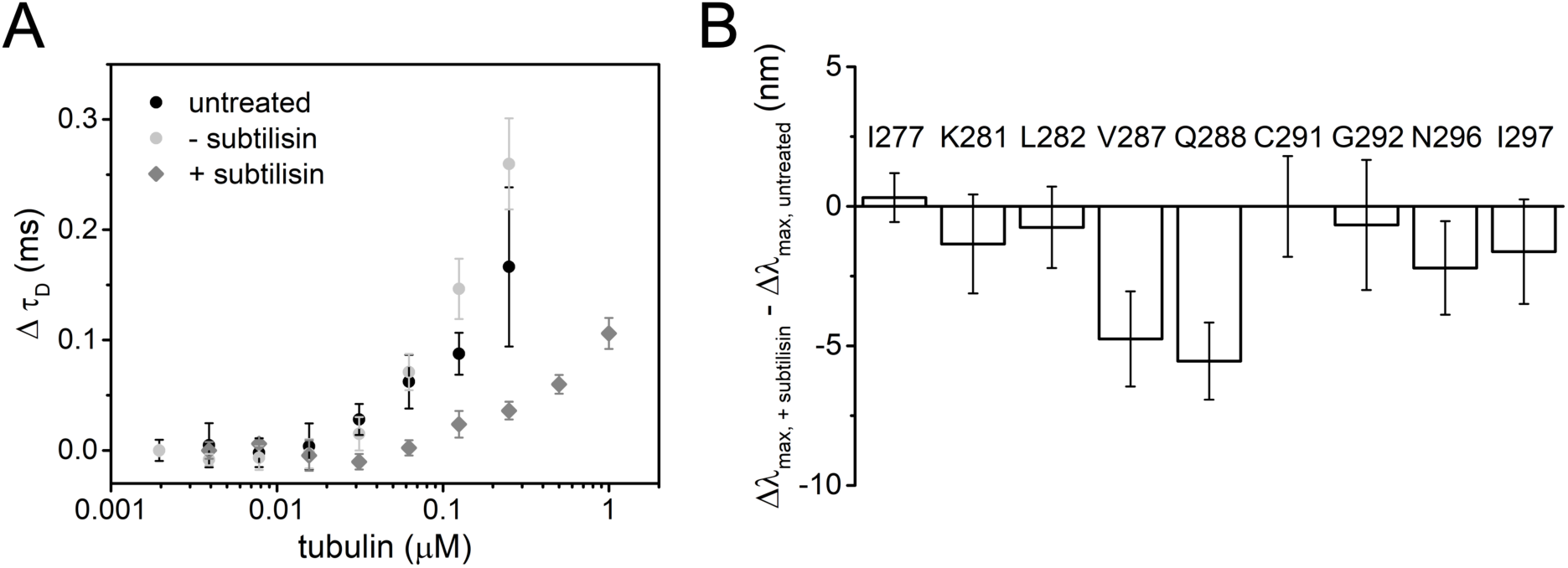
Disordered tails of tubulin enhance tau binding but minimally impact R2 structure. (A) Δτ_D_ of 20 nM tau_A488_ upon binding to tailless or intact tubulin. Data points are mean ± SD, n=3. See also Figure S3. (B) Δλ_max_ of tau_acrylodan_ upon binding to 4 μM tailless tubulin relative to 2 μM intact tubulin. At these respective tubulin concentrations, saturation of tau:tubulin binding is achieved without formation of crosslinked complexes. Data points are mean ± SD, n≥3.

When bound to tailless tubulin, the Δλ_max_ for tau_acrylodan_ were either not significantly different from, or only slightly smaller than, those measured when bound to intact tubulin for most of the positions tested (Figure 2B). This indicates that the environment of most R2 residues is not significantly impacted by the presence or absence of the tails. The largest decreases in acrylodan shifts were observed at V287 and Q288 (Figure 2B). Although from the EM model, these residues are distant from the originating sites of the tubulin tails (Figure 1C), the tails nevertheless appear to influence the stability of the interaction of these two residues with the α-tubulin surface. N296 and I297 also showed moderate decreases in acrylodan shifts; these residues are found on the surface of the adjacent β-tubulin in the cryo-EM structure (Figure 1C and 2B). As our FCS measurements suggest that tau binds fewer tailless tubulin dimers than intact tubulin dimers (Figure S3B), these residues may be impacted by the absence of tubulin at the adjacent binding site in R3.

### Hierarchy in tubulin occupancies in MTBR

Overall the magnitude of the blue shifts in the acrylodan spectra for the R2 probes examined here are lower than those we observed previously in R3 (Figure 1D) (Li et al., 2015). To gain insight into these differences, we labeled tau at two different equivalent positions within each repeat (Figure 3A): K281, selected because it is located at the intradimer tubulin interface near the center of the stretch of residues resolved by cryo-EM and has the largest peak shifts in our measurements (Figure 1C); and D283, which is completely conserved across different repeats and points into the solvent in the cryo-EM structure (Figure 1C). The Δλ_max_ were measured as a function of tubulin concentration in order to assess the apparent affinity of binding by each repeat (Figure 3B and C). The R3 site showed the largest peak shifts at all concentrations of tubulin (Figure 3B – E). The trend in Δλ_max_ for both positions was R3>R2>R4≈R1 (Figure 3D and E), in good agreement the C291-equivalent residues tested in our previous work (Li et al., 2015). While the magnitude of the Δλ_max_ differed for the different repeats, the biphasic behavior observed with K281 was present in all of the repeats in both positions tested, with saturating tubulin concentrations comparable to those measured by FCS (Figure 1B and 3B and C). Normalization of the peak shift curves to their values at 10 μM tubulin was plotted to compare the apparent affinity for individual repeats. Comparison of the D283 equivalent positions reveal equivalent affinities for each repeat (Figure 3C), whereas the K281 equivalent positions show that R3 (acrylodan at P312) has a slightly higher affinity for tubulin compared to the rest of the repeats (Figure 3B). Interestingly, the concentration dependence of P312 is similar to that of the D283 equivalent positions (Figure 3C). This suggests that mutating M250, K281 and K343 had caused a slight decrease in affinity, ∼2-fold (Figure 3), consistent with the speculation that these residues may interact with the tubulin tails (Kellogg et al., 2018). The overall similarities in affinities of the different repeats suggests that the differences in the magnitudes of Δλmax likely reflects occupancy of tubulin binding sites in each repeat.

**Figure 3.**
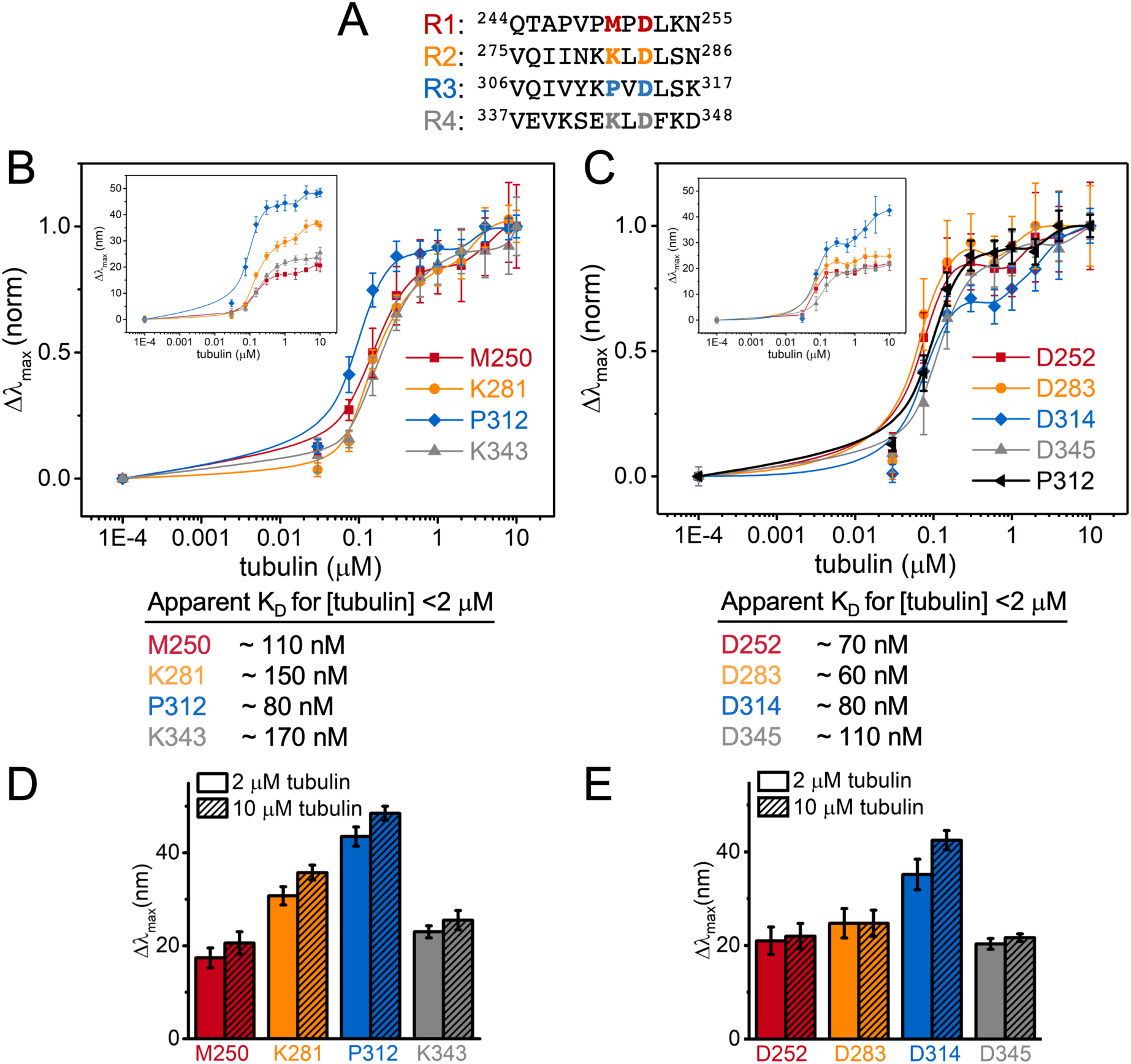
Hierarchy in tubulin occupancies in MTBR. (A) Equivalent residues tested highlighted in red, orange, blue and grey for R1 – R4 respectively. Normalized Δλ_max_ of tau_acrylodan_ at (B) K281 and (C) D283-equivalent positions upon binding to tubulin. Tau only measurements were plotted at 1E-4 μM tubulin for visualization. Data points are mean ± SD, n≥3. Insets are the raw Δλ_max_. Approximate K_D_s from binding to [tubulin] <2 μM are reported below. (D and E) Bar graph for Δλ_max_ with 2 μM or 10 μM tubulin for comparison.

## Discussion

While isolated tau has been described as disordered since it was first purified and characterized (Cleveland et al., 1977; Woody et al., 1983), the details of its tubulin- or MT-bound structure have been elusive. This is in part due the fact that low-affinity binding sites for tubulin and MT are distributed throughout tau (Butner and Kirschner, 1991; Mandelkow et al., 1995), resulting in a highly dynamic interaction. With soluble tubulin, the interaction is even more complex, as variable population of the tubulin binding sites results in a heterogenous distribution of tau:tubulin complexes (Li and Rhoades, 2017). Here we use an environmentally sensitive fluorophore, in combination with a published structural model (Kellogg et al., 2018), to interrogate the structural features of tau in tau:tubulin complexes and gain insight into tau-mediated polymerization of tubulin.

Our acrylodan data for the MTBR of tau bound to soluble tubulin is largely consistent with the recent structural model for MT-bound tau (Figure 1) (Kellogg et al., 2018). This conclusion is derived from the Δλ_max_ profile of tau_acrylodan_ at different sites within R2, which showed good correlation between emission blue-shift and surface accessibility calculated from the EM model (Figure 1E). Interestingly, the hydrophobic residues probed – I277, L282, L284 and V287 – all of which interface with tubulin were found to have lower acrylodan emission blue-shifts than neighboring polar or charged residues – K281, S285 and Q288 – all of which are oriented away from the tubulin surface (Figure 1C). The smaller than expected Δλ_max_ are likely due to a reduction in the shift for the hydrophobic, tubulin-interfacing residues due to the probe’s interaction with tubulin surface charges, as described in the Results. We first encountered this apparent anomaly in the relationship between residue hydrophobicity and Δλ_max_ of acrylodan-labeled R3 (Li et al., 2015) (Figure S2D), which we are able to resolve in this study through the context gained from the EM structure.

There are weak binding sites for tubulin distributed throughout tau’s MTBR. As a consequence, the fraction of binding sites occupied is highly dependent upon the tubulin concentration (Figure S1). There are two ways to interpret differences in Δλ_max_. The first possibility is that differences in the Δλ_max_ of equivalent positions across different repeats reflect differences in tubulin occupancy, i.e. a rank order of residency: R3>R2>R4≈R1 (Figure 3). In this case, the larger magnitude Δλ_max_ for R2 and R3 results from greater occupancy of these binding sites with respect to R4 or R1. This suggests that tubulin binding between repeats may not be independent, i.e. the occupancy of R3 increases the probability of occupancy of R2 (Figure 4, Model 1). The larger shift in R3 relative to the other repeats may indicate a preference for R3 when coupled with its neighboring repeats. These results are consistent with prior work from our lab with soluble tubulin which found that mutations at equivalent residues in R2 and R3 had differential effects on binding to tubulin, with the impact of the R3 mutations being much larger (Elbaum-Garfinkle et al., 2014). This is in contrast to that observed for MTs, where R1 and R2 are thought to dominate the interaction (Butner and Kirschner, 1991; Goode et al., 1997; Goode and Feinstein, 1994). Alternately, differences in Δλ_max_ could be simply due to differences in local environment when bound to tubulin, i.e. equivalent positions in different repeats do not experience equivalent interactions with tubulin (Figure 4, Model 2). Considering the persistent rank order present in all two equivalent positions tested here, as well as in our prior work (Li et al., 2015), we speculate that the coupled occupancy model (Model 1) is the dominant contributor, and that the large differences in the Δλ_max_ of R3 positions relative to the other repeats may also reflect differences in the local environment of R3 (Model 2). While tau’s role in stabilizing microtubules is well-established, much less is known about the physiological significance of its interactions with soluble tubulin. Coordinated binding of tubulin dimers in adjacent repeats, may, for example, enhance tau’s ability to nucleate MT polymerization (Li and Rhoades, 2017). Alternatively, tau may differentiate between binding to soluble tubulin and the MT-lattice through preferential interactions of different repeats. Localizing preferred binding domains for soluble tubulin and MTs to different repeats also prompts consideration as to whether tau can bind simultaneously to both.

**Figure 4.**
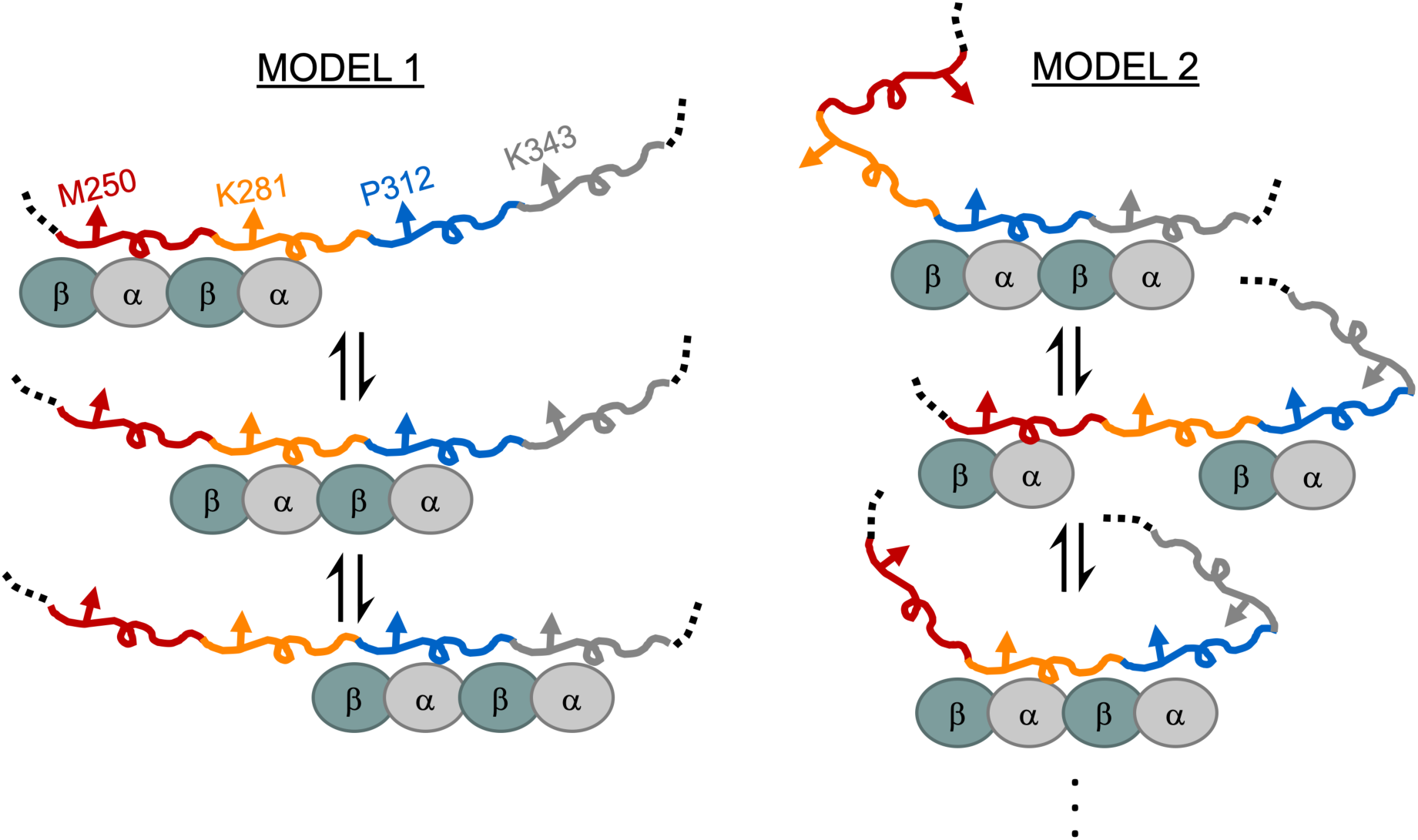
Proposed models for heterogeneity in tau:tubulin complexes. Tau’s MTBR is drawn in segments colored as in Figure 3 and tubulin dimers are depicted as cyan and grey ovals. Model 1 depicts cooperative binding of neighboring repeats which causes the enrichment of R2 and R3 occupancy over R1 and R4. Model 2 depicts uncoupled binding of tubulin to different repeats and structural changes of tau generates differences in environment of the probed position.

Interestingly, the disordered tubulin tails have a significant impact on both tau binding affinity and on the size and heterogeneity of tau:tubulin complexes, but do not have a large impact on tau’s tubulin-bound conformation (Figure 2). The reduction in formation of large tau:tubulin assemblies suggest critical roles of tubulin tails in forming fuzzy complexes that are important for tau-mediated polymerization (Li and Rhoades, 2017). An interaction between tau and disordered tubulin tails on MTs has been reported previously (Chau et al., 1998; Di Maïo et al., 2014; Serrano et al., 1985) with the tail-binding site mapping to R1 and the inter-repeat region linking R1 and R2 on tau (Chau et al., 1998). Tau has also been shown to directly bind to synthetic peptides corresponding to the C-terminal tails of both α- and β-tubulin (Chau et al., 1998; Lefèvre et al., 2011; Maccioni et al., 1989) and that the isolated tails can compete with MTs for tau binding (Devred et al., 2004). Our data demonstrates that while the structure of tubulin-bound R2 is not greatly altered by the removal of tubulin tails (Figure 2B), residues such as K281 that were shown to be solvent facing in the EM structure (Kellogg et al., 2018) contribute weakly to binding (Figure 3B). This suggests that the interaction between the tails and R2 is likely to be transient and dynamic, and that the loss of affinity seen with tailless tubulin may be due to interactions with regions of tau outside the MTBR, such as the proline-rich region. Further elucidation of tubulin and MT binding sites in tau domains flanking the MTBR will provide critical insights into how they influence tau function.

In summary, we have used site-specific acrylodan fluorescence to probe structural features of tau:tubulin complexes. Our data reveal that the structural similarities between tubulin- and MT-bound tau. Moreover, we find evidence of hierarchical binding sites within the MTBR which point to a possible mechanism of differentiation between tubulin and MT interactions relevant to tau function. Our study underscores the potential of site-specific fluorescence, in conjunction with an established structural model, to provide critical insights in structure-function relationships of disordered, dynamic systems. This combination of approaches may be a viable strategy for studying other challenging and dynamic systems that stymie more traditional structural characterization.

## Acknowledgements

We thank D. Chenoweth for use of his fluorometer and the Department of Chemistry Mass Spectrometry Facility for use of the Bruker Ultraflex III. Sedimentation velocity measurements and analysis were performed at the Johnson Foundation Structural Biology and Biophysics Core Facility at the University of Pennsylvania’s Perelman School of Medicine. This work is funded by NIH NINDS AG053951.

## Author Contributions

Conceptualization, H.Y.J.F. and E.R.; Investigation, H.Y.J.F, K.M, J.R. and K.G.; Writing – Original Draft, H.Y.J.F., K.M., K.G. and E.R.; Writing – Review & Editing, H.Y.J.F., and K.G. and E.R.; Funding Acquisition, E.R.

## Declaration of Interests

The authors declare no competing interests.

## STAR Methods

### Contact for Reagents and Resources Sharing

Further information and requests from resources and reagents should be directed to and will be fulfilled by the Lead Contact, Elizabeth Rhoades.

### Experimental Model and Subject Details

*E. coli* strain BL21(DE3) was used for expression of recombinant proteins for *in vitro* studies, and growth was in LB medium.

Tubulin was purified from fresh brains from young cows obtained from a local slaughterhouse.

## Method Details

### Recombinant tau purification and labeling

Tau (residues 149 – 395) was cloned into a pET vector with an N-terminal HisX6 tag and a TEV cleavage site. For FCS measurements, natural C322 was mutated to serine and the remaining C291 was labeled with Alexa Fluor 488 maleimide (tau_A488_). For ensemble fluorescence measurements, both C291 and C322 were mutated to serine and individual sites were mutated to cysteine for labeling with acrylodan. Tau constructs were expressed in BL21(DE3) cells in 1 liter growths in LB media at OD_600_ ≈0.5 – 0.6 by induction with 1 mM IPTG for 4 hrs at 37°C. Cells were pelleted and resuspended in Buffer A (50 mM Tris pH 8.0, 500 mM NaCl, 10 mM imidazole) supplemented with 1 mM PMSF and a cOmplete protease inhibitor cocktail tablet. The lysate was then flash-frozen in liquid nitrogen and stored at −80°C until use. Lysozyme (1 mg/mL) was added to the thawed lysate immediately prior to sonication, and the sonicated lysates were then clarified by centrifugation at 20,000xg for 30 mins at 4°C. The supernatant was filtered through a 0.45 μm syringe filter and then incubated with Ni-NTA beads for 1 hr at 4°C with gentle rocking. The Ni-NTA beads were then washed with Buffer A and tau was eluted with Buffer B (50 mM Tris pH 8.0, 500 mM sodium chloride, 400 mM imidazole). Eluted protein was buffer exchanged into Buffer C (25 mM Tris pH 8.0, 150 mM NaCl and 1 mM DTT) using 10k MWCO Amicon concentrators prior to incubation with TEV protease at room temperature for 2 – 2.5 hrs. The protein samples were passed through Ni-NTA beads to remove cleaved HisX6 tag and TEV protease (also His-tagged) and further purified by size-exclusion chromatography over a Superdex 200 HiLoad 16/600 column in Buffer D (25 mM Tris pH 8.0, 100 mM NaCl, 1 mM EDTA and 0.5 mM TCEP).

For labeling, the proteins were concentrated to ∼300 μM (ε_λ=280nm_= 4470) and treated with 1 mM DTT for 30 mins at room temperature to reduce the cysteines. The proteins were then buffer exchanged into Buffer E (20 mM Tris pH 7.4, 50 mM NaCl) with 6 M guanidine HCl using HiTrap desalting columns. For Alexa Fluor 488 labeling, the C5 maleimide dye in a DMSO stock was added in 2 – 4 times molar excess and incubated at room temperature for 30 mins followed by overnight incubation at 4°C. For acrylodan labeling, DMSO was added to the protein sample to 10% final concentration prior to the addition of dye to 4 times molar excess and incubated for 4 hrs at room temperature. In both cases unconjugated dye and guanidine HCl was removed from the protein samples by two rounds of buffer exchange into Buffer E using concentrators and a final desalting column step. Labeled proteins were aliquoted, snap frozen and stored at −80°C until use. Complete labeling of tau proteins were confirmed by MALDI-TOF mass spectrometry using a Bruker Ultraflex III instrument. Sinapic acid dissolved in 50:50 acetonitrile:water (v/v) and 0.1% TFA was used as a matrix.

### Tubulin purification and tail cleavage

Tubulin was purified from young bovine brains as described (Castoldi and Popov, 2003), with the GTP concentration in the first polymerization step increased to 1 mM. Purified tubulin was stored at aliquots at −80 °C. For FCS and acrylodan fluorescence measurements, tubulin was quickly thawed, centrifuged at 100,000xg for 6 mins at 4°C and buffer exchanged using Bio-spin 6 columns into Buffer F (20 mM potassium phosphate pH 7.4, 20 mM KCl, 1 mM MgCl_2_, 0.5 mM EGTA and 1 mM DTT) and used within 1 hr after thawing.

Tailless tubulin was generated by modification of a published protocol (Knipling et al., 1999). Briefly, tubulin was quickly thawed and buffer exchanged into 1/10X Buffer G (0.1 M MES pH 6.9, 1 mM MgCl_2_ and 1 mM EGTA) using Bio-spin 6 columns and diluted to 50 μM. GTP was added to 1 mM and the mixture was incubated at room temperature for 5 mins. Subtilisin was then added to 16.8 μg/mL (1:300 w/w) and the reaction was incubated for 45 mins at room temperature. PMSF (0.1 mM) was added to quench the reaction and the mixture was incubated on ice for 10 mins before centrifugation at 110,000xg for 20 mins. Supernatant containing tailless tubulin was buffer exchanged back into Buffer G for storage at −80°C. Complete cleavage of the tails was confirmed by MALDI-TOF. All subtilisin-treated tubulin had a loss of ∼1.8-2 kDa when compared to intact tubulin, consistent with the loss of tails from both α and β subunits. Prior to use, aliquots of tailless tubulin were clarified and buffer exchanged as described above.

### Fluorescence correlation spectroscopy (FCS)

FCS measurements were performed on a lab-built instrument based around an inverted Olympus IX-71 microscope (Melo et al., 2017; Nath et al., 2010). Briefly, a 488 nm laser was focused into sample with a 60x 1.2 NA water immersion objective. The emitted fluorescence was collected through the objective and separated from the excitation light via Z488RDC and 500LP filters. The collected emission was focused into a 50 μm optical aperture fiber directly coupled to a photon counting module. A digital correlator was used to calculate autocorrelation curves. For measurements mirroring conditions of fluorimeter assays using 300 nM tau, the laser power was adjusted to be ∼1 μW at the back aperture of the objective to avoid high photon count rates on the photon diodes. For all other measurements, 20 nM tau and 5.0 μW laser power was used. Measurements were made in borosilicate coverglass chambers that were pre-treated with PEG-PLL to reduce protein adsorption (Huang et al., 2001). Tau_A488_ was mixed with varying concentrations of tubulin in Buffer F and incubated for 5 mins at room temperature prior to measurements. For each experiment, 40 – 50 autocorrelation curves of 10 secs were collected from each sample. All experiments were performed in triplicate and on different days with different protein aliquots.

### Sedimentation Velocity Analytical Ultracentrifugation (SV-AUC)

SV-AUC experiments were performed at 20°C with a Beckman-Coulter XL-A analytical ultracentrifuge and a TiAn60 rotor with two-channel charcoal-filled epon centerpieces and quartz windows. Samples were analyzed at either 280 nm or 485/490 nm for the detection of total protein or tau_A488_, respectively. Concentrations were selected to obtain best signal in both wavelengths: tubulin, 8.7 μM; unlabeled tau, 90 μM; tau_A488_,16 μM; tau_A488_ and tubulin mixture, 3.5 μM and 10.5 μM, respectively. Samples were buffer exchanged and prepared into Buffer F. Complete sedimentation velocity profiles were recorded every 30 seconds at 40,000 rpm.

### Acrylodan fluorescence

Fluorescence emission of tau_acrylodan_ was measured using a Horiba Fluorolog-3 fluorometer in a Quartz cuvette. Excitation was set at 390 nm and emission was scanned from 400 to 600 nm. Both excitation and emission slit widths were set at 2 nm. The cuvette holder was held at 20°C. 300 nM tau_acrylodan_ was mixed with varying tubulin concentrations in Buffer F and incubated in an eppendorf tube for 5 mins prior to measurement. Either 3 or 4 independent measurements were made for each tau construct using different tubulin aliquots across multiple days.

### Fluorescence anisotropy

Fluorescence emission of tau_acrylodan_ (labelled at I277 and K281) and tau_A488_ (labelled at position T149) were measured on a JASCO-8300 fluorometer with temperature control accessory ETC-815. Tau_acrylodan_ was excited at 390 nm and scanned from 400 to 600 nm, and tau_488_ was excited at 470 nm and scanned from 480 to 650 nm. Both excitation and emission slits were set to 2.5 nm. Clarified tubulin was buffer exchanged into Buffer F and incubated at varying concentrations with 300 nM tau for 5 minutes prior to measurement.

## Quantification and Statistical Analysis

### Analysis of FCS data

Autocorrelation curves were fitted using Python in Anaconda-Spyder with a single component fit using the following equation:

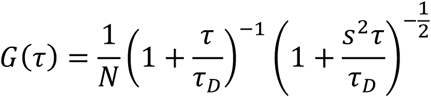

where *G*(*τ*) is the autocorrelation as the function of time *τ, N* is the average number of fluorescent molecules, *τ*_*D*_ is the average diffusion time of the molecules and *s* is the ratio of radial to axial dimensions of the focal volume, determined using Alexa 488 as a reference standard and fixed to 0.23/0.24 for all data fitting. All autocorrelation curves were fitted with the above equation and filtered by goodness-of-fit and Anderson-Darling tests for normal and log-normal distributions as described previously (McKibben and Rhoades, 2019). This approach removes autocorrelation curves arising from infrequent but large, bright tau assemblies that otherwise skew analysis of the majority species (Li and Rhoades, 2017).The autocorrelation curves from filtered data sets from each measurement were then averaged and fitted with equation above. Differences in diffusion times were calculated by subtracting diffusion times with that of tau only (Figure 1B and S3B) or lowest tubulin concentration used (Figure 2A). Data points presented in Figure 1B, 2A and S3B are means and standard deviations of triplicate measurements. For counts per molecule analyses (CPM) in Figure S1B and S3D, data obtained from different days were combined prior to filtering (total of 150 – 200) and diffusion time and CPM of individual curves after filtering were reported with means of the whole population indicated. CPM for each trace were calculated by dividing its average intensity by *N* and was normalized with the average CPM obtained from the tau only measurement. Data were plotted using Origin.

### Analysis of SV-AUC data

The solvent density (ρ = 1.0022 g/mL), and viscosity (η = 0.01013 poise) were derived from chemical composition by the program SEDNTERP (Laue et al., 1992). A standard partial-specific volume (Ū) value of 0.73 cm^3^/g was used in all analysis. Data were assessed in two complementary ways. First, data were analyzed using the program *DCDT+* version 1.02 (Dr. John S. Philo, Thousand Oaks, CA) was used, which employs the time-derivative algorithm (Philo, 2006). This software was used to obtain apparent g(s*) distribution plots of sedimenting species (i.e. uncorrected for diffusion) (Stafford 1992). The g(s*) data were derived from scans using a point spacing of 0.5 S. Secondly, data were also fit using the Lamm equation as implemented in the program SEDFIT (Schuck, 2000). After optimizing meniscus position and fitting limits, the sedimentation coefficient (S) and best-fit frictional ratio (*f/f*_*0*_) was determined by iterative least squares analysis. Data were plotted using Origin.

### Analysis of acrylodan emission peak shifts

Each emission trace was background subtracted with buffer or tubulin only measurements. Maximum emission wavelength was determined using Origin peak search algorithm using 1^st^ derivative method with smoothing. For each labeling position, measurements were made of tau in the absence and presence of tubulin and the mean emission maximum of tau in the absence of tubulin was used to calculate the shift in the emission peak, Δλ_max_. The emission maximum for tau labeled with acrylodan in the absence of tubulin was relatively insensitive to position, with peaks ∼522 nm (Figure S2B). The emission peaks shifts are reported as means and standard deviations. The error is propagated in the calculations for changes in peak shifts. In Figure 3B and C, normalization was performed using the measured peak shift at 10 μM tubulin and the standard deviation was error-propagated. Data were plotted using Origin and the accompanying structure figure (Figure 1C) was generated using Pymol.

### Analysis of fluorescence anisotropy

Spectra were integrated to calculate total fluorescence intensities in order to account for peak shifts in the acrylodan upon binding. The anisotropy (r) was calculated as follows:

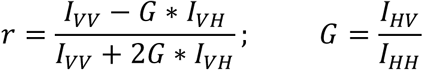

where *I*_*VV*_, *I*_*VH*_, *I*_*HV*_, and *I*_*HH*_ are the integrated intensities for each respective excitation and emission polarizer position, V=vertical polarization, H=horizontal polarization. Mean r and standard deviations were reported from 2 to 3 independent measurements. The normalized change in anisotropy (r_norm_) was calculated by subtracting the tau-only anisotropy from the tau + tubulin anisotropy and normalizing the data from 0 to 1. Error bars on this data were not error propagated and are shown to illustrate variability of the independent measurements. Data were plotted using Origin.

