## Supplemental Information for "Structural characterization of tau in fuzzy tau:tubulin complexes"

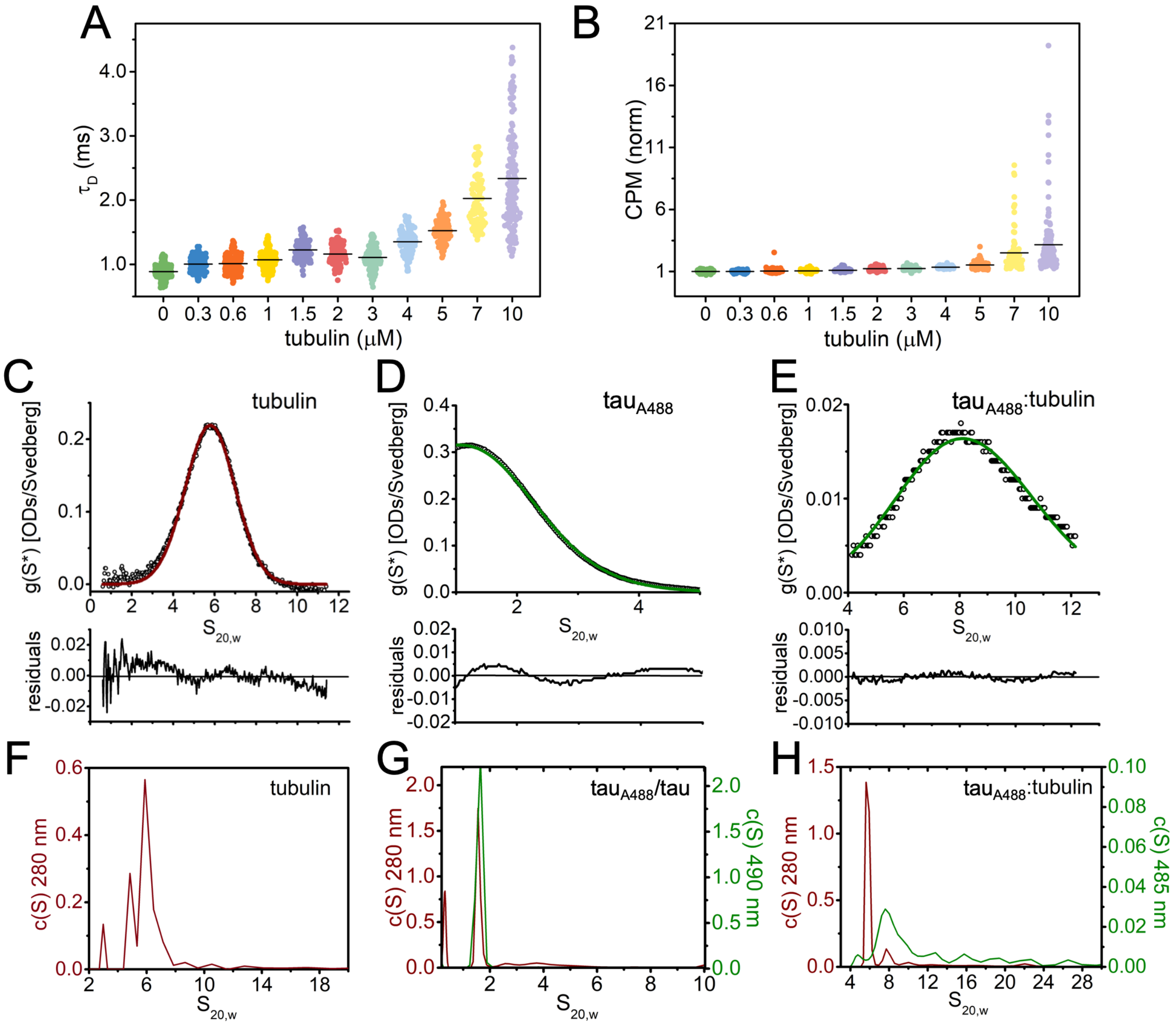


**Figure S1. Related to Figure 1. Analysis of tau:tubulin binding.** (A) $\tau_D$ and (B) normalized CPM from analysis of individual autocorrelation curves are plotted with black lines indicating the mean. (C – E) Sedimentation coefficient distribution, g(s*), was calculated from the time-derivative algorithm of the concentration profile as implemented in the program *DCDT*+ (Philo, 2006; Stafford, 1992) for tubulin (C, 280 nm), $tau_{A488}$ (D, 490 nm), and a 1:3 $tau_{A488}$:tubulin (E, 485 nm) using SV-AUC measurements. Upper panels are g(s*) plots of primary data (grey circles) fitted to a one-species Gaussian function (solid lines) and lower panels are fitting residuals. $S_{20,w}$ values obtained are 5.9 ± 0.006 S for tubulin, 1.6 ± 0.015 S for $tau_{488}$, and 8.3 ± 0.010 S for $tau_{A488}$:tubulin respectively (mean ± SD). (F – H) c(S) distributions derived from the fitting of the Lamm equation to the same experimental data collected, as implemented in the program SEDFIT (Schuck, 2000). Double-Y plots are shown, with c(S) distributions derived from 280 nm detection in brown and those from 490/485 nm detection in green. $S_{20,w}$ = 8.3 S suggests that tau:tubulin complexes at 1:3 molar ratio contain on average more than one but less than two tubulin dimers. This molar ratio is lower than that of the first saturation in FCS curves (1:6.67) due to limiting tubulin concentrations (already at 10.5 μM), and suggests that complexes at the first saturation likely contain two tubulin dimers.

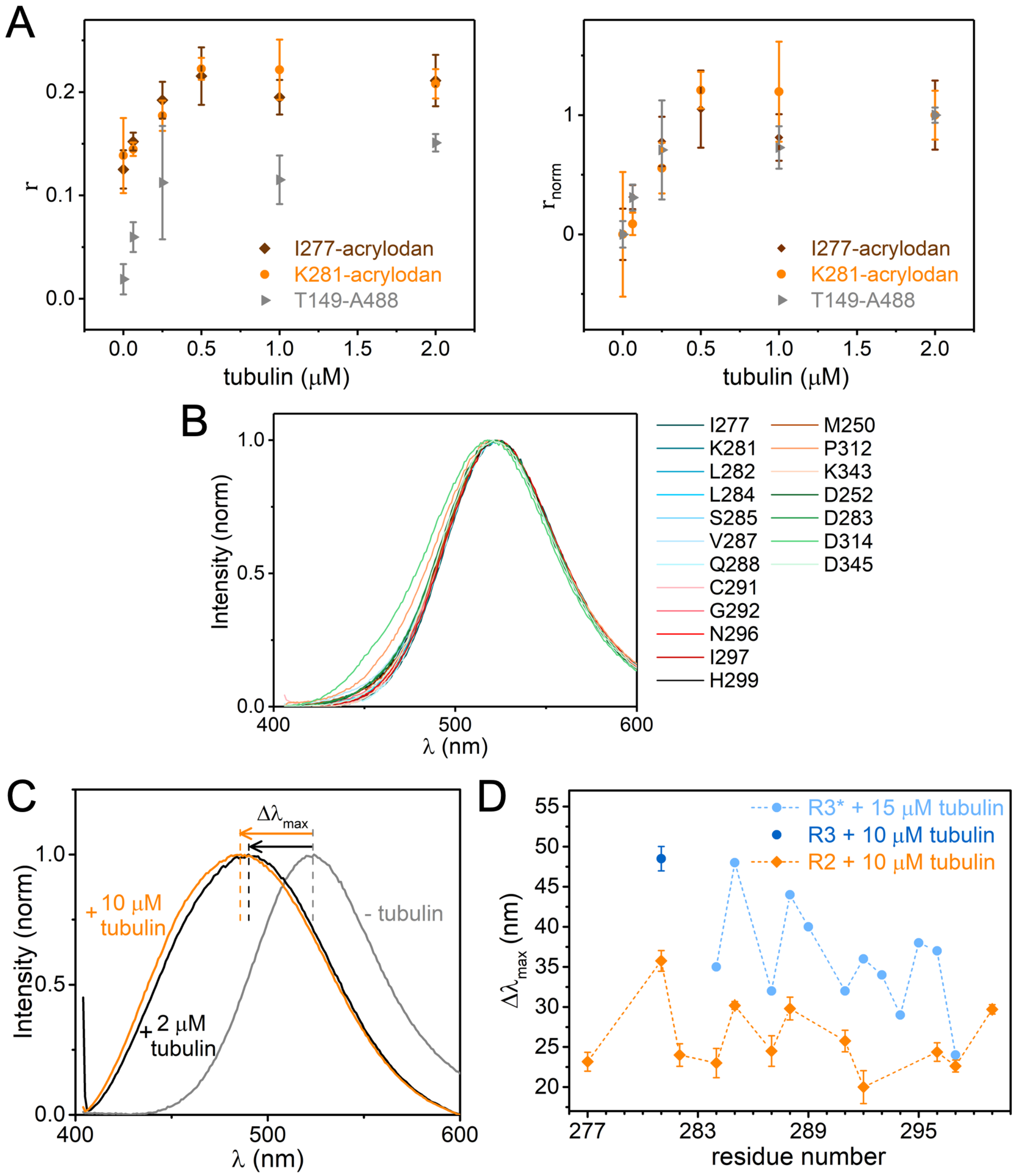


**Figure S2. Related to Figure 1. Tau labeled with acrylodan binding to tubulin** (A) Anisotropy (r and $r_{norm}$) of tau labelled with acrylodan at I277 and K281 or with Alexa Fluor 488 at T149 titrated with tubulin. Data points are mean ± SD, n=2 – 3. Error bars in normalized plot on the left are not error-propagated. (B) Emission spectra of $tau_{acrylodan}$ in all positions tested in this study. Intensities are normalized to peak maximum for comparison. Peak emission was not affected by location of acrylodan conjugation. (C) Representative plots of emission blue-shifting ($\Delta\lambda_{max}$) of $tau_{acrylodan}$ (at K281) in the presence of 2 μM and 10 μM tubulin. (D) $\Delta\lambda_{max}$ for residues tested in R2 (data from Figure 1D in orange) and R3-equivalent positions (data from Figure 3B in blue and those previously tested in Li et al., 2015 in light blue) are plotted for comparison. Data is replotted from Li et al., 2015 with permission. *Tau construct used in this study spans residues 198 – 372, as compared to the 149 – 395 construct used in this study.

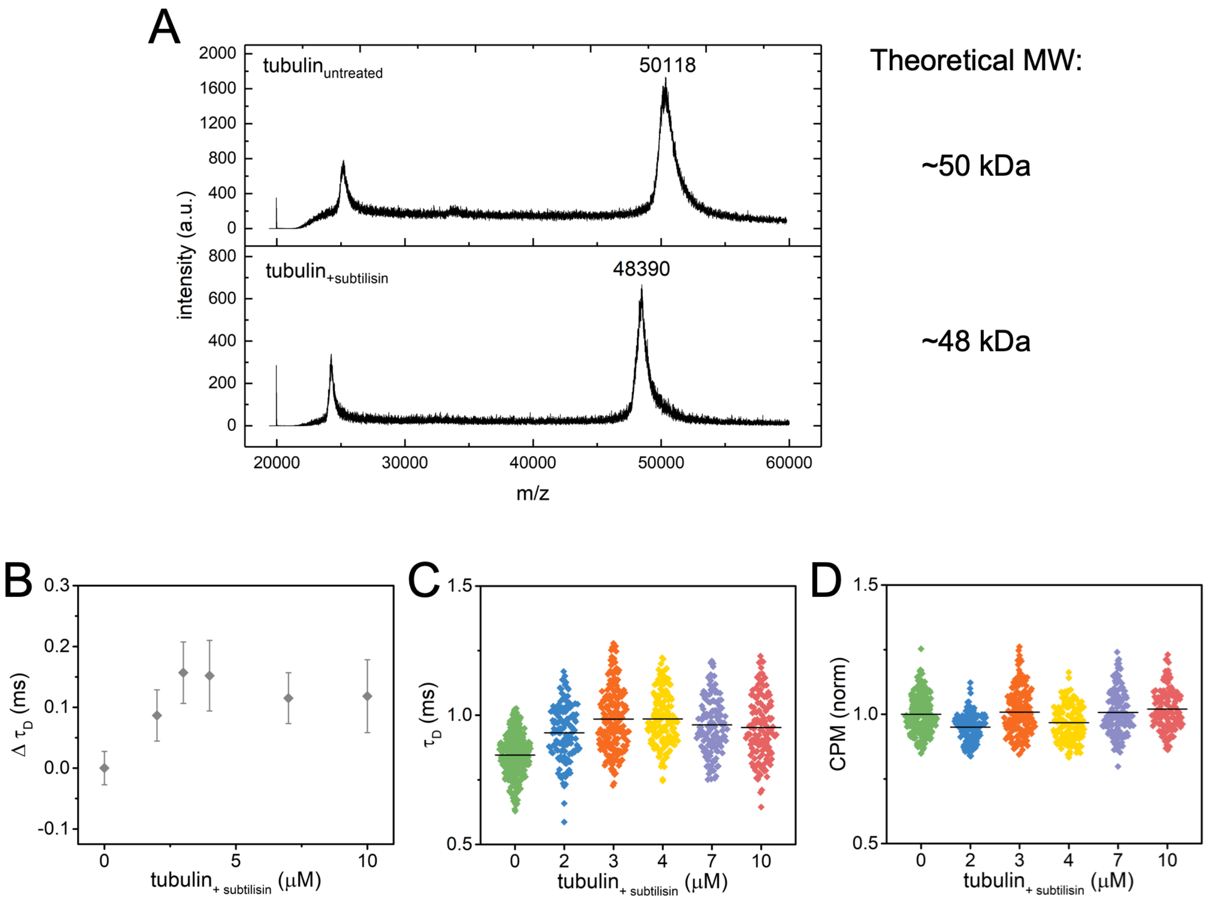


**Figure S3. Related to Figure 2. Tau binding to tailless tubulin** (A) MALDI-TOF spectra of intact (upper) and tailless (lower) tubulin. The difference in mass between intact and tailless tubulin is <2 kDa, consistent with the removal the disordered tails from both tubulin monomers (Knipling et al., 1999). (B) Changes in $\tau_D$ of 300 nM $tau_{A488}$ upon binding tailless tubulin measured by FCS. Data points are mean ± SD, n=3. (C) $\tau_D$ and (D) normalized CPM from analysis of individual autocorrelation curves are plotted black lines indicating the means.

**KEY RESOURCES TABLE**

| REAGENT or RESOURCE | SOURCE | IDENTIFIER |
|---|---|---|
| Bacterial and Virus Strains | | |
| *E. coli* strain BL21-CodonPlus(DE3)-RIL | Agilent Technologies | 230240 |
| Biological Samples | | |
| Young cow brains | Marco Farms, PA | N/A |
| Chemicals, Peptides, and Recombinant Proteins | | |
| cOmplete protease inhibitor cocktail tablet | Sigma-Aldrich | 11873580001 |
| IPTG | Gold Bio | I2481 |
| Ampicilin | Gold Bio | A301 |
| Lysozyme | Sigma-Aldrich | L6876 |
| PMSF | Gold Bio | P470 |
| TEV protease | Produced in-house | N/A |
| DTT | Thermo Fisher | BP172-5 |
| TCEP-HCl | Gold Bio | TCEP |
| Alexa Fluor 488 C5 Malemide | Thermo Fisher | A10254 |
| Acrylodan | Thermo Fisher | A433 |
| Sinapic Acid | Sigma-Aldrich | D7927 |
| Proteinase, bacterial (Subtilisin) | Sigma-Aldrich | P8038 |
| GTP | Sigma-Aldrich | G8877 |
| PEG-PLL | Produced in-house | N/A |
| Recombinant DNA | | |
| pET vector with Hisx6-TEV-hTau(149 – 395, C291S C322S + single cysteine mutations at various sites for labeling) | This paper | N/A |
| Software and Algorithms | | |
| Digital correlator | Correlator.com | Flex03lq-12win7 |
| Spyder | Anaconda | https://anaconda.org/anaconda/spyder |
| Origin 2016 | Origin Lab | https://www.originlab.com/2016 |
| SEDNTERP | Laue et al., 1992 | http://bitcwiki.sr.unh.edu/index.php/Main_Page |
| DCDT+ version 1.02 | Philo, 2006 | http://www.jphilo.mailway.com/dcdt+.htm |
| SEDFIT | Schuck, 2000 | http://www.analyticalultracentrifugation.com/default.htm |
| Pymol | Schrodinger LLC | https://pymol.org/ |
| Other | | |
| Ni-NTA Agarose | Qiagen | 30230 |
| 0.45μm syringe filter | Milipore | SLHVM33RS |
| 10kDa MWCO Amicon concentrators | Milipore | UFC901024 |
| Superdex 200 HiLoad 16/600 | GE Healthcare | 28989335 |
| HiTrap desalting columns | GE Healthcare | 17140801 |
| Micro Bio-spin 6 columns | Bio-rad | 7326221 |
| Sapphire 488-50 laser | Coherent | N/A |
| Photon Counting Module | Excelitas | SPCM-AQR-14-FC |

| | | |
|---|---|---|
| 8-well Nunc Lab-Tek #1.0 coverglass chambers | Thermo-Fisher | 155411 |
